# Factors associated with cholera outbreaks, Nairobi County, July 2017: a case control study

**DOI:** 10.1101/719641

**Authors:** Valerian Mwenda, Alexis Niyomwungere, Elvis Oyugi, Jane Githuku, Mark Obonyo, Zeinab Gura

## Abstract

**Background:** Cholera affects 1.3-4 million people globally and causes 21000-143,000 deaths annually. Nairobi County in Kenya reported cholera cases since April 2017. We investigated to identify associated factors and institute control measures.

**Methods:** We reviewed the line-list of patients admitted at the Kenyatta National referral Hospital, Nairobi and performed descriptive epidemiology. We carried out a frequency-matched case control study, using facility-based cases and community controls. We defined a case as acute onset of watery diarrhoea of at least >3 stools/24hours with or without vomiting in person of any age, admitted in Kenyatta National Hospital as from July 1^st^, 2017. We calculated odds ratios and their respective 95% confidence intervals. We also took water samples at water reservoirs, distribution and consumer points, and made observation on hygiene and sanitation conditions in the community.

**Results:** We reviewed 71 line-listed cases; median age 30 years (range 2-86 years); 45 (63%) were male. First case was admitted on 14^th^ April 2017. Culture was performed on 44 cases, 30 (68%) was positive for *Vibrio cholerae*, biotype El-Tor, serotype Ogawa. There were 2 deaths (case fatality ratio 2.8%). Age-group ≥25 years was most affected. Drinking unchlorinated water (aOR 14.57, 95% CI 4.44-47.83), eating in public places (aOR 9.45, 95% CI 3.07-29.12) sourcing water from non-Nairobi city water company source (aOR 4.92, 95% CI 1.56-15.52) and having drank untreated water in the previous week before the outbreak (aOR 3.21, 95% CI 1.12-9.24) were independently associated with being a case in the outbreak. Out of 28 water samples, 4 (14%) had >180 coliforms/100mls; all were at consumer points.

**Conclusion:** Poor water quality and sanitation were responsible for this outbreak. We recommended adequate, clean water supply to unplanned settlements in Nairobi County, as well education of residents on water treatment at the household level.

**Author summary:** Cholera, a disease causing outbreaks in areas with low standards of hygiene and sanitation has afflicted humans for millennia. It is caused by a bacterium, *Vibrio Cholerae*, transmitted mainly through water contaminated by faecal matter. The resultant disease is acute watery diarrhoea, which causes death rapidly due to dehydration and shock. Virtually brought under control in the developed world due to improvements in hygiene, the disease still ravages many communities in low and middle income countries, as well as regions affected by conflict or natural disasters. In outbreak situations, rapid response in water treatment, sanitation improvement and setting up of cholera treatment centres for rehydration therapy reduces impact and saves lives. Long-term control can only be achieved through sustainable improvements in sanitation and standards of living. Case control studies in outbreak situations provide quick, actionable information to public health specialists during outbreak response. This study provides a cholera outbreak investigation in an urban informal settlement setting; the approach reported here can guide in outbreak investigations and response in similar settings globally.

## Introduction

Cholera is an enteric infection caused by toxigenic *Vibrio cholerae* serogroup O1 and O139 [1]. Cholera is endemic in more than 50 countries globally and causes large epidemics in countries or regions facing complex emergencies including conflict, natural disasters like flooding or drought or massive displacement of persons [2]. It is estimated that every year, 1.3 billion people are at risk, 1.3 to 4 million get infected and 21000 to 143000 deaths occur globally [3]. Up to 90% of infected people continue shedding the bacterium up to 14 days without or with mild symptoms making cholera outbreaks difficult to control [4].

Most cholera outbreaks occur in areas with inadequate supply of potable water and poor sanitation facilities, especially during rainy seasons [5]. Other factors identified as risk factors in previous epidemiological studies done in Kenya include lack of knowledge about cholera, proximity to a large water body, living in a refugee camp and eating food outside the home [6–10]. A cross-district analysis of cholera occurrence also identified open defaecation as a risk factor [11].

Increase in acute watery diarrhoea cases, confirmed to be cholera by culture, was reported in Nairobi County from April 2017 through the Integrated Disease Surveillance and Reporting (IDSR) system. Sporadic cases were reported in May and early June but an upsurge was observed in late June and July 2017. The epicentre of the outbreak was unplanned settlements from the Eastern suburbs of the city; however, cases were also reported in high end hotels and restaurants. Most of the cases were admitted at Kenyatta National Referral Hospital, since an ongoing nurses’ industrial action had paralyzed tier one, two and three facilities [12]. The Ministry of Health (MoH) deployed a team comprising of the Disease Surveillance and Response Unit (DSRU), the Kenya Field Epidemiology and Laboratory Training Programme (FELTP), Nairobi County Department of Health, National Public Health Laboratory (NPHL) and the Ministry of Water (MoW) to identify the associated factors and institute control.

## Methods

### Investigation setting

This investigation was carried out during July 24-28, 2017 in various locations in Nairobi County including the Kenyatta National Hospital (KNH) where most patients were admitted for care and five sub-counties: Embakasi South, Embakasi East, Embakasi West, Mathare and Starehe (Figure 1). These were the sub-counties from which the cases originated from. Nairobi County has an estimated population of 3.5 million residents as of 2016 and 6.5 million with the suburbs included. Approximately 2.5 million of these live in unplanned settlements [13]. The city is divided into 17 sub-Counties, with 12 of them having reported cholera cases during the period Since May 2017.

**Figure 1:**
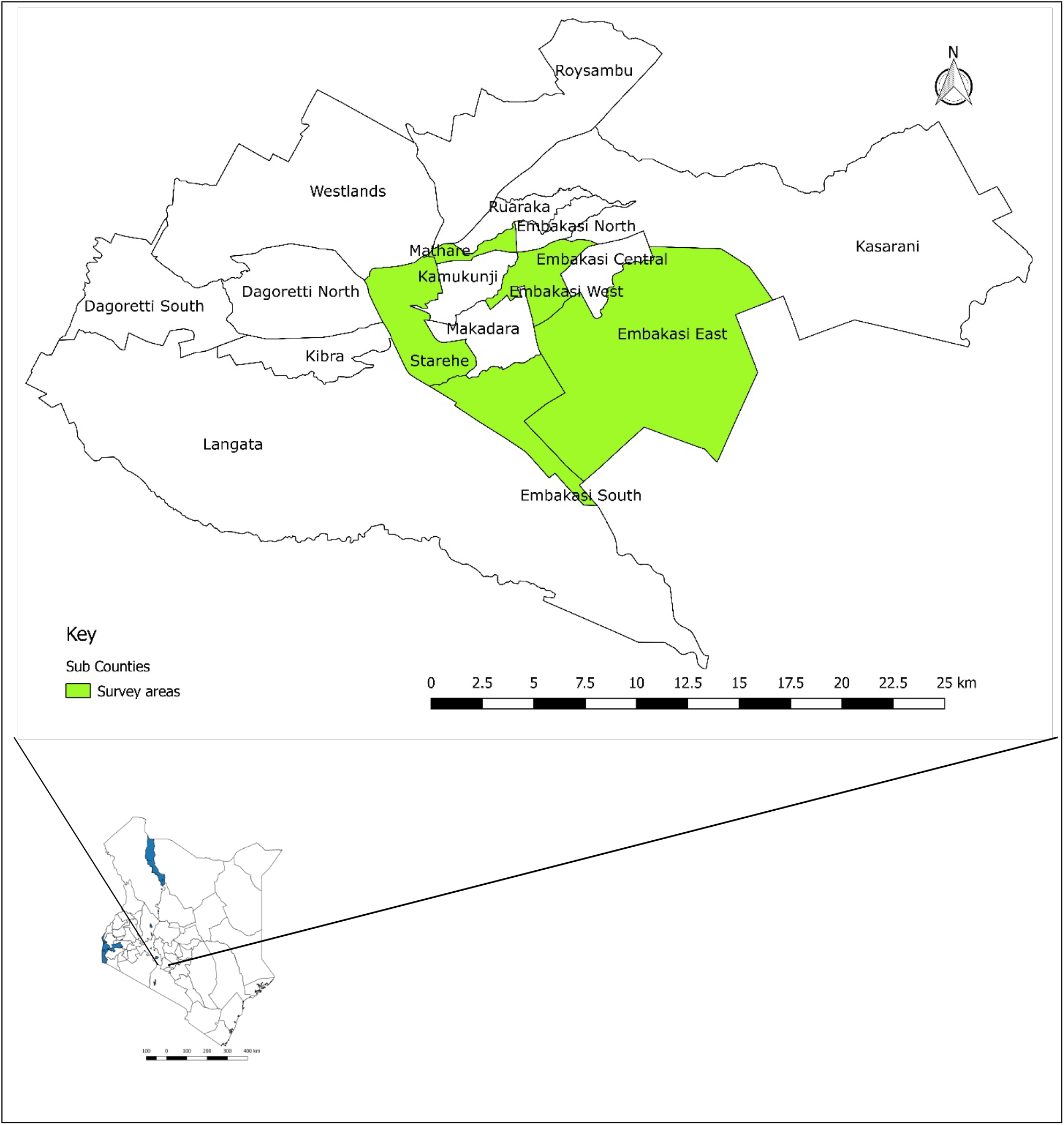
Map of Nairobi County, showing the study sites (Sub-counties affected by the Cholera outbreak) highlighted in green. (Maps produced using QGIS, a free and open source Geographic Information System, available at https://qgis.org/en/site/).

### Retrospective review

We reviewed the cholera line list from KNH up to July 16, 2017. The variables contained in the line list included Sub County of origin, residence (village/estate), age, sex, date of onset/admission/discharge, laboratory status and patient outcome. We conducted descriptive analysis of the updated and consolidated line list, describing the outbreak in terms of place of origin, time of symptoms onset and personal attributes like age and sex.

### Case control

We then carried out a frequency matched case control study using facility cases from KNH and community controls from the most affected sub-counties. Cases were matched to controls using age groups (2-4, 5-14, 15-24, >24) years on a ratio of one case to two controls. Two years is the cut-off for cholera case definition in areas with active outbreaks as per the IDSR guidelines [14].

### Case definition

We included both probable and confirmed cases in our investigation. A probable case was defined as acute onset of watery diarrhoea of at least >3 stools/24hours with or without vomiting in a person ≥2 years, admitted in Kenyatta National Hospital as from July 1^st^, 2017. A confirmed case was culture positive.

Controls were defined as absence of diarrhoea in the preceding 30 days in any randomly selected person of the same age group as a particular case and came from the same Nairobi Sub-counties as cases. We administered verbal screening for symptom and admission questions to any control prior to enrolment into the study.

#### Inclusion criteria

Those who met the case/control definition and consented verbally to the study.

## Sample Size Calculation

### Assumptions

We made the following assumptions while calculating the sample size for the study: Power 80%, 21.6% prevalence of exposure for hand washing before meals among controls [15] to detect a least an exposure odds ratio of 3.0, desired two-sided confidence intervals of 95% and a case: control ratio of 1:2. A minimum sample size of 132 (44 cases and 88 controls) was calculated using the Fleiss method [16].

### Selection of cases

Cases appearing in the consolidated KNH line list since July 1^st^, 2017 and still admitted in the hospital were eligible for inclusion in the study. The admission register in cholera treatment ward served as the sampling frame. We used simple random sampling to select the cases, and administered a structured questionnaire through face to face interview. Verbal consent was sought from the cases and legal guardians in case of a minor.

### Selection of Controls

For each case we selected two population controls, distributed into five sub-counties. The sub-counties were selected on the basis of the number of cases reported, with the three with the most number of cases and two with the least number of cases. We visited the selected sub-counties, and used the administrative offices of the Sub-counties at as our starting point. Spinning a bottle to choose direction of proceeding, we selected every fifth household for selection of controls. In areas where the direction was interrupted by infrastructural installations like roads or industrial complexes, we spun the bottle a repeat time, to cover the expanse of the unplanned settlements. The resident Community Health Volunteers guided us on the demarcation of various households, due to the high population density in the settlements. We administered a structured questionnaire similar to the one for cases on the demographics and risk factor sections, but without clinical details section. Before the interview, controls were screened for cholera-like symptom history, including diarrhoea, vomiting, abdominal pains, and cholera case in the household, in the previous one month.

### Environmental testing

We collected water samples for testing from the Nairobi city water company pipeline system and households, using sterile containers. For the water company system, the testing was done at the water treatment sites/ reservoirs, during piping and selected consumer points. Samples were also collected from conveniently selected control households, from the affected sub-counties, as well as from some public establishments like schools in the affected sub-counties. Analysis of the samples was undertaken at the NPHL. General bacteriological analysis through the most probable number (MPN) was used to estimate faecal coliform count. Levels of residual chlorine were measured using automated colorimetry and physical conductivity tests done [17].

### Data management

Data obtained was entered into a computer database, cleaned and analysed. Measures of central tendency and dispersion for continuous variables and proportions for categorical variables were calculated. We calculated odds of various exposures among the cases and controls and corresponding odds ratios (OR). Factors with a P-value of ≤0.15 [18] at bivariate analysis were included in unconditional logistic regression model, using the forward selection approach. A confidence interval excluding the null value of OR significant in the final model. During logistic regression, the matching variable (age group) was maintained in the model till the end of the analysis.

### Ethical considerations

Informed consent was obtained orally from all study participants and recorded in the interview questionnaire; written consent was difficult because the study was conducted during emergency outbreak response. Confidentiality of the information from the participants was maintained at all times. We did not collect any personal identification information, the questionnaires were kept in a locked cabinet during data entry and the computer database created protected by a password, accessible only to the principal investigator. This being a public health emergency response, we did not seek approval of the investigation from an independent research and ethics committee. However, approval was sought from the Ministry of Health and permission to conduct the investigation from the County Government of Nairobi Department of Health.

## Results

### Retrospective review

The consolidated line list had 71 cases as at July 16, 2017. The median age was 30 years (Inter-quartile range 12.5); 45 (63%) were males. A total of 44 cases had culture done; 30 (68%) were positive. There were 2 deaths, case fatality rate 2.8%. Age group above 25 years was the most affected (Table 1).The three Embakasi Sub-Counties (East, West and South) had the highest number of cases at 25 (36%). The first cholera case was admitted to KNH on April 14, 2017; sporadic cases were admitted in May 2017. Several peaks were observed in late June and early July 2017 (Figure 2).

**Table 1:** Attack rates for different age groups and sex, cholera outbreak in Nairobi County, July 2017

| Age | Age group | Number of cases | Total population | Attack rate (per 100,000 population) |
| --- | --- | --- | --- | --- |
|  | <4 years | 2 | 407988 | 0.49 |
|  | 5-14 | 6 | 564906 | 1.06 |
|  | 15-24 | 7 | 753209 | 0.93 |
|  | ≥25 | 56 | 1412266 | 4.00 |
| Sex | Male | 45 | 1605230 | 2.80 |
|  | Female | 26 | 1533139 | 1.70 |

**Figure 2:**
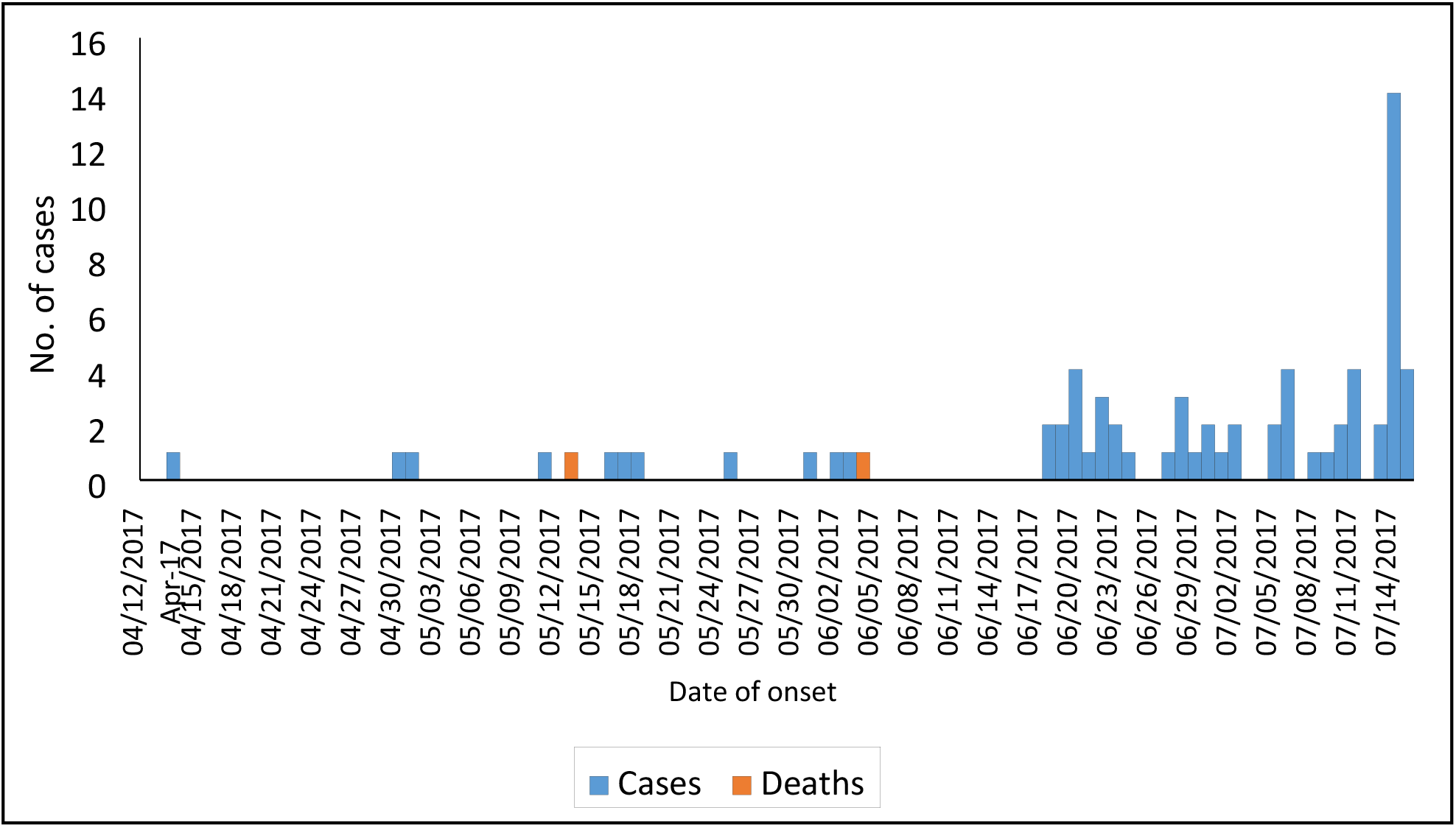
Epidemic curve, Cholera patients at Kenyatta National Hospital, July 2017 (N=71)

### Case control findings

Mean age was 30.9 years±9.0 years for cases and 28.8 years±9.0 years for controls; 31 (71%) of cases and 30 (33%) of controls were male. Thirty-five cases (80%) and 55 (60%) controls had completed secondary education (Table 2). Most of the cases had presented with watery diarrhoea (98%) and vomiting (80%). The median hospitalization period for the cases was 3 days, range 2-5 days. Only 19 (43%) of the cases sought care within 6 hours initial symptoms.

**Table 2:**
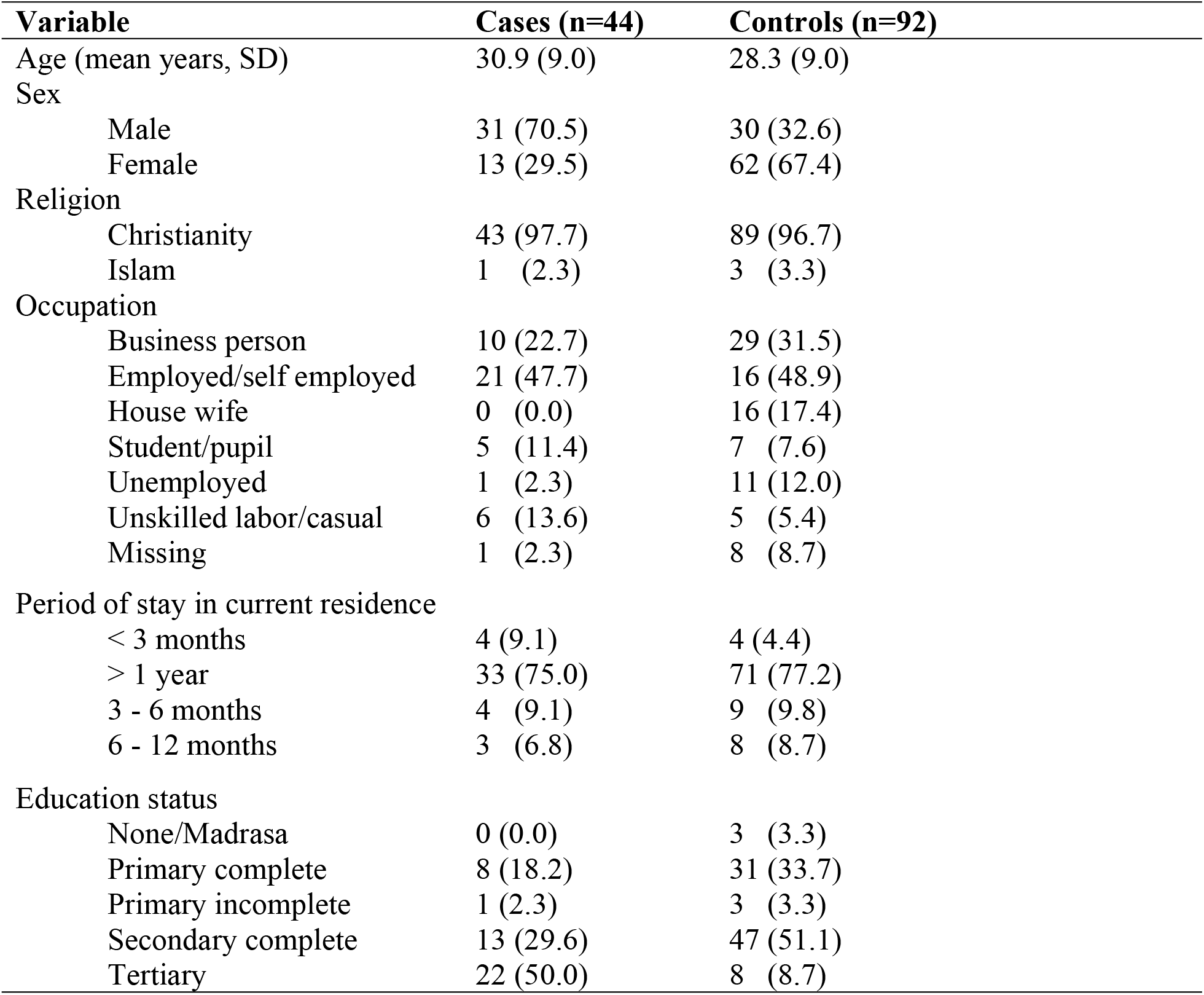
Sociodemographic characteristics of the study participants

### Bivariate analysis

Cases had 10 times higher odds of using unchlorinated water (OR 10.31, 95% CI 3.96-26.89), nine times higher odds of taking meals at public eating places (OR 8.97, 95% CI 3.52-22.30) and not washing hands after toilet (OR 8.91, 95% CI 2.46-39.62) at bivariate analysis. (Table 3).

**Table 3:** Bivariate and multivariate analysis of factors associated with cholera outbreak, Nairobi County, July 2017

| Exposure variable | Cases n (%) | Controls n (%) | Crude OR (95 % CI) | aOR |
| --- | --- | --- | --- | --- |
| Not Treating drinking water |  |  |  |  |
| Yes | 21 (47.7) | 19(20.7) | 3.51 (1.61-7.63) | NA |
| No | 23 (52.3) | 73 (79.3) | Ref |  |
| Drinking water from Borehole |  |  |  |  |
| Yes | 6 (13.6) | 1 (1.1) | 14.37 (1.63-667.91) | NA |
| No | 38 (86.4) | 91 (98.9) | Ref |  |
| Drinking water from non-municipal supplies |  |  |  |  |
| Yes | 20 (45.5) | 12 (13.0) | 5.56 (2.38-12.98) | 4.92 (1.56-15.52) |
| No | 24 (54.5) | 80 (87.0) | Ref |  |
| Storing drinking water in uncovered container |  |  |  |  |
| Yes | 10 (22.7) | 4 (4.4) | 6.47 (1.70-29.74) | NA |
| No | 34 (77.3) | 88 (95.6) | Ref |  |
| Drinking untreated water in previous week |  |  |  |  |
| Yes | 32 (74.4) | 39 (52.0) | 2.69 (1.18-6.10) | 3.21 (1.12-9.24) |
| No | 11 (25.6) | 36 (48.0) | Ref |  |
| Not chlorinating water before drinking |  |  |  |  |
| Yes | 38 (86.4) | 35 (38.0) | 10.31 (3.96-26.89) | 14.57 (4.44-47.83) |
| No | 6 (13.6) | 57 (62.0) | Ref |  |
| Not Washing hands before eating |  |  |  |  |
| Yes | 10 (22.7) | 3 (3.3) | 8.53 (2.0-50.22) | NA |
| No | 34 (77.3) | 87 (96.7) | Ref |  |
| Not Washing hands after visit to toilet |  |  |  |  |
| Yes | 13 (29.6) | 4 (4.5) | 8.91 (2.46-39.62) | NA |
| No | 31 (70.4) | 85 (95.5) | Ref |  |
| Eating away from home |  |  |  |  |
| Yes | 37 (84.1) | 31 (37.3) | 8.87 (3.52-22.30) | 9.45 (3.07-29.12) |
| No | 7 (15.9) | 52 (62.7) | Ref |  |

*Ref: reference group, NA: not included in logistic regression, OR: odds ratio, aOR: adjusted odds ratio*

### Multivariate analysis

After controlling for confounders, drinking unchlorinated water (aOR 14.57, 95% CI 4.44-47.83), taking meals at public eating places (aOR 9.45, 95% CI 3.07-29.12) and sourcing drinking water from non-city water company sources (aOR 4.92, 95% CI 1.56-15.52) were independently associated with being a case in the outbreak (Table 3).

### Water testing and environmental findings

Out of 28 water samples, four (14%) had >180 coliforms/100ml of water; from a restaurant, secondary school kitchen, residential apartments and a public water point. Of the 17 samples that underwent conductivity and residual chlorine testing, all had between 0.4-1.2 ppm (recommended range 0.2-5.0) of residual chlorine and 74.3-87.0µS/cm (recommend range 0-800) on conductivity levels.

## Discussion

Our investigation found use of untreated drinking water and taking meals away from home as risk factors in this outbreak in a major urban setting. Association with taking meals at public eating establishments was also a finding during the investigation of protracted outbreaks in Kenya in 2014-2016 [10] and is important in prevention of future urban food-borne outbreaks. Majority of urban residents take at least one meal daily away from their households highlighting importance of safe public eating places. Some of the water samples at consumer points were contaminated.

Cholera outbreaks have affected various counties in Kenya since December 2014 [19]. Water and sanitation is a major factor in cholera transmission in most outbreaks globally [20]. The current outbreak affected relatively younger individuals and males more than females. Males are likely to consume different types of meals regarded as ‘high risk’ outside the home during occupational ventures, hence predisposing them to higher risk of contracting food borne diseases [21]. The case fatality of 3% for the patients admitted at the referral facility (KNH) is unexpectedly high, with CFR expected to be less than 1% if proper treatment is instituted promptly [22]. This can be explained by several contextual realities affecting both the Nairobi County and National public health system at the time. Nurses were in the middle of an industrial action, therefore all the public dispensaries and health centres were non-functional. Since these usually serve as the avenues for setting up Cholera Treatment Centres (CTC), the resultant effect was delayed proper management and late referral to the national referral hospital. Peaks of cases in June and July were as a result of outbreaks during mass gathering of people on various occasions [23].

On risk factor analysis, using untreated water and taking meals in public eating places were noted as important exposures associated with being a case in the outbreak. Since the epicentre of this outbreak was from informal settlements in the city, contaminated water as well as unhygienic eating places, especially serving casual labourers in the adjacent industries were likely avenues for disease transmission. The affected sub counties boarder the Nairobi city industrial complex, where many young men work as casual labourers and eat from food vendors, a recognized ecological risk factor for the disease [24]. Majority of the residents were getting water that is illegally piped in unhygienic environment, especially in open sewer trenches. Failure to undertake domestic treatment of this water before use is therefore a major point of exposure to many waterborne diseases.

The water sampled from the city water company at holding reservoirs and distribution networks had adequate residual chlorination and conduction levels. Therefore contamination of the four samples likely occurred downstream during transmission to households. This could be caused by illegal connections into the distribution network. Contaminated water is the main vehicle of cholera transmission worldwide and offers opportunities for disease control [25].

We found several gaps in response to the outbreak. First, the initial cases were not fully investigated till the peaks in June and July occurred; this was a lost opportunity since promptly investigating and instituting control measures reduces extent, scope and possibility of propagation of outbreaks [26]. No cholera treatment centres (CTCs) were set up in the affected Sub-counties; patients had to be ferried to the referral facility likely aiding disease transmission. The best practice would have been to treat cholera at the sites of the outbreak. The nurses’ strike reduced the effectiveness of the response, and likely contributed to the protracted course of the outbreak. Provision of water to most informal settlements in Nairobi is inadequate; water vendors and illegal connections fill in the gap but expose residents to unpotable water for human consumption.

Cholera control and prevention can be achieved in various ways. Of these, water, sanitation and hygiene improvement are the most effective, with water treatment preventing up to 90% of water borne diseases [27]. Cholera vaccination has only been effective in outbreak situations when offered together with provision of safe water and improving environmental hygiene [28]. Primary prevention in form of sanitizing the environment and provision of safe water are also effective against cholera and other water and food borne diseases.

Our investigation had limitations. The sampling of the water was not random, as advised by the WHO [29] therefore may not be representative to the water in use in the settlements from which the cases and controls came from. We also did not manage to tests the water samples for *Cholerae vibrio*, nor did we associate the contaminated water samples with the exact origin of the cases.

### Public health action

We supported the Nairobi County Department of Health (CDoH) in setting up and operationalizing CTCs in the two most affected sub-counties, as well as health education and provision of water treatment supplies. We also trained the Community Health Volunteers on first aid and quick response when assisting cholera victims as well as the Sub-County disease surveillance coordinators on proper data capture, management and timely transmission to the DSRU. The communities were sensitized on cholera symptoms, importance of prompt seeking of care if the disease is suspected, importance of water, sanitation and hygiene (WASH) practices and various methods of water treatment.

### Conclusion

We confirmed that this cholera outbreak, with epicentres in several informal settlements in Nairobi, was associated with taking untreated water and eating meals at public eating places. Long term control requires investment in improving clean water supply to the informal settlements, in adequate amounts, throughout the year.

## Acknowledgements

We wish to thank the Nairobi County Department of health, Sub-county disease surveillance officers and community health assistants for their support during this study.

## Supporting information

S1: Article submission cover letter.

S2: Filled STROBE Checklist for a case control study.

## References

1. Harris JB, LaRocque RC, Qadri F, Ryan ET, Calderwood SB: Cholera. The Lancet, 379(9835):2466–2476.

2. Jutla A, Khan R, Colwell R: Natural Disasters and Cholera Outbreaks: Current Understanding and Future Outlook. Current environmental health reports 2017, 4(1):99–107.

3. Ali M, Nelson AR, Lopez AL, Sack DA: Updated global burden of cholera in endemic countries. PLoS neglected tropical diseases 2015, 9(6):e0003832.

4. Barzilay EJ, Schaad N, Magloire R, Mung KS, Boncy J, Dahourou GA, Mintz ED, Steenland MW, Vertefeuille JF, Tappero JW: Cholera surveillance during the Haiti epidemic—the first 2 years. New England Journal of Medicine 2013, 368(7):599–609.

5. Jutla A, Whitcombe E, Hasan N, Haley B, Akanda A, Huq A, Alam M, Sack RB, Colwell R: Environmental Factors Influencing Epidemic Cholera. The American Journal of Tropical Medicine and Hygiene 2013, 89(3):597–607.

6. Mutonga D, Langat D, Mwangi D, Tonui J, Njeru M, Abade A, Irura Z, Njeru I, Dahlke M: National surveillance data on the epidemiology of cholera in Kenya, 1997-2010. The Journal of infectious diseases 2013, 208(suppl_1):S55–S61.

7. Mohamed AA, Oundo J, Kariuki SM, Boga HI, Sharif SK, Akhwale W, Omolo J, Amwayi AS, Mutonga D, Kareko D: Molecular epidemiology of geographically dispersed Vibrio cholerae, Kenya, January 2009-May 2010. Emerging infectious diseases 2012, 18(6):925.

8. Onyango D, Karambu S, Abade A, Amwayi S, Omolo J: High case fatality cholera outbreak in Western Kenya, August 2010. Pan African Medical Journal 2013, 15(1).

9. Stoltzfus JD, Carter JY, Akpinar-Elci M, Matu M, Kimotho V, Giganti MJ, Langat D, Elci OC: Interaction between climatic, environmental, and demographic factors on cholera outbreaks in Kenya. Infectious diseases of poverty 2014, 3(1):37.

10. George G: Notes from the field: ongoing cholera outbreak-Kenya, 2014-2016. MMWR Morbidity and mortality weekly report 2016, 65.

11. Cowman G, Otipo S, Njeru I, Achia T, Thirumurthy H, Bartram J, Kioko J: Factors associated with cholera in Kenya, 2008-2013. Pan African Medical Journal 2017, 28(101).

12. Lancet: Kenya’s nurses strike takes its toll on health-care system. The Lancet, 389(10087):2350.

13. World Population Review: Kenyan population. Available at http://worldpopulationreview.com/countries/kenya-population/. Last accessed December 4, 2017.

14. Kenya Integrated Disease Surveillance and Response Guidelines. Available at http://guidelines.health.go.ke:8000/media/Standard_Case_Definitions_for_Priority_Diseases_in_Kenya-_Integ.pdf. Last accessed April 12, 2018. In.

15. Hutin Y, Luby S, Paquet C: A large cholera outbreak in Kano City, Nigeria: the importance of hand washing with soap and the danger of street-vended water. Journal of Water and Health 2003, 1(1):45–52.

16. Ibrahim MA: The Case-control Study Consensus and Controversy, vol. 32: Elsevier; 2014.

17. Organization WH: Guidelines for drinking-water quality: first addendum to the fourth edition. 2017.

18. Bursac Z, Gauss CH, Williams DK, Hosmer DW: Purposeful selection of variables in logistic regression. Source code for biology and medicine 2008, 3(1):17.

19. WHO. Emergency preparedness and response. Cholera in Kenya. Available at http://www.who.int/csr/don/21-july-2017-cholera-kenya/en/. Last accessed December 4, 2017.

20. Wright J, Gundry S, Conroy R: Household drinking water in developing countries: a systematic review of microbiological contamination between source and point-of-use. Tropical medicine & international health 2004, 9(1):106–117.

21. Shiferaw B, Verrill L, Booth H, Zansky SM, Norton DM, Crim S, Henao OL: Sex-based differences in food consumption: Foodborne Diseases Active Surveillance Network (FoodNet) population survey, 2006-2007. Clinical infectious diseases 2012, 54(suppl_5):S453–S457.

22. WHO. Prevention and control of cholera outbreaks. Available at http://www.who.int/cholera/technical/prevention/control/en/. Last accessed April 2, 2018

23. Ministry of Health, Kenya. Cholera response, 2017. Available at http://www.health.go.ke/2017/07/government-close-two-hotels-to-contain-cholera-outbreak/ Last accessed January 15, 2018.

24. Hunter P: Waterborne disease: epidemiology and ecology: John Wiley & Sons; 1997.

25. Taylor DL, Kahawita TM, Cairncross S, Ensink JH: The impact of water, sanitation and hygiene interventions to control cholera: a systematic review. PLoS one 2015, 10(8):e0135676.

26. Mwambi P, Mufunda J, Lupili M, Bangwe K, Bwalya F, Mazaba M: Timely Response and Containment of 2016 Cholera Outbreak in Northern Zambia. Medical Journal of Zambia 2016, 43(2):64–69.

27. Fewtrell L, Kaufmann RB, Kay D, Enanoria W, Haller L, Colford JM: Water, sanitation, and hygiene interventions to reduce diarrhoea in less developed countries: a systematic review and meta-analysis. The Lancet infectious diseases 2005, 5(1):42–52.

28. Clemens J, Holmgren J: When, how, and where can oral cholera vaccines be used to interrupt cholera outbreaks? In: Cholera Outbreaks. edn.: Springer; 2013: 231–258.

29. Organization WH: Guidelines for drinking-water quality [electronic resource]: incorporating 1st and 2nd addenda, vol. 1, Recommendations. 2008.

